# Mesoscopic Fluorescence Imaging of Light-Triggered Chemotherapeutic Release in Cancer Spheroid Models

**DOI:** 10.64898/2026.02.08.704660

**Authors:** Elias Kluiszo, Rasel Ahmmed, Berna Aliu, Semra Aygun-Sunar, Matthew Willadsen, Hilliard L Kutscher, Jonathan F. Lovell, Ulas Sunar

## Abstract

Peritoneal micrometastases (micromets) remain a major barrier to durable cytoreduction in ovarian and other intra-abdominal cancers, because lesions can be difficult to visualize and are often resistant to systemic therapy. Liposomal doxorubicin (Dox) improves pharmacokinetics but can be limited by slow intratumoral release. Porphyrin-phospholipid (PoP) liposomes enable near-infrared light–triggered release of Dox (chemophototherapy (CPT)), creating an opportunity for intraoperative, fluorescence-guided treatment planning and monitoring. Here, we evaluate a laparoscopic fluorescence imaging platform for quantifying light-triggered drug delivery in 2D monolayers and 3D spheroid cluster models. Dox fluorescence increased linearly with administered LC-Dox-PoP concentration in both SCC2095sc and SKOV-3 cultures (R^2^ = 0.97-0.98 in 2D; R^2^ = 0.98 in spheroid clusters over 1–9 µg/mL). Laparoscope-derived fluorescence measurements agreed with standard well-plate reader measurements (R^2^ = 0.89-0.96). Porphyrin fluorescence provided stronger, complementary contrast for localizing spheroid constructs and decreased after activation light exposure, consistent with photobleaching during triggered release. Together, these results support a quantitative imaging framework for fluorescence-guided monitoring of light-triggered liposomal drug release, with potential to inform individualized CPT dosimetry for peritoneal micrometastases. These findings in SCC2095sc (oral squamous cell carcinoma) additionally suggest relevance of fluorescence-guided CPT for head and neck/oral cancer, where localized post-resection adjuvant treatment may improve control of residual disease.

## 1. Introduction

Peritoneal micrometastases (micromets) represent significant hurdles in the successful treatment of intra-abdominal malignancies, specifically ovarian cancer, which remains one of the most lethal gynecological diseases due to high recurrence rates [1–3]. These micromets are often too small to be detected by conventional imaging techniques such as computed tomography (CT), magnetic resonance imaging (MRI), positron emission tomography (PET), and ultrasound, which often demonstrate lower sensitivity than reassessment surgeries [4–8]. Consequently, even after surgical resection and adjuvant chemotherapy, up to 60% of patients may still harbor occult disseminated disease [1,2,4,5,7,9]. Thus, there is an urgent need for sensitive intraoperative imaging methods that allow surgeons to visualize and selectively treat these lesions in real-time while avoiding damage to healthy tissue.

An emerging solution lies in chemophototherapy (CPT), which combines light-activated drug release with photodynamic effects [10–15]. While traditional systemic chemotherapy often leads to significant toxic side effects [3,7,13,15–18], these liposomal nanocarriers improve the biodistribution and efficacy of the payload [18,19]. In our approach, doxorubicin (Dox) is encapsulated within porphyrin-phospholipid (PoP) liposomes, improving biodistribution while enabling spatially confined, near-infrared light–triggered release (~660 nm) at the illuminated site [10,11,16,17]. The lipid envelope protects the drug from degradation and facilitates passive accumulation at tumor sites via the enhanced permeability and retention (EPR) effect [6,20,21]. In parallel, photodynamic therapy (PDT) leverages light, a photosensitizing agent, and molecular oxygen to generate cytotoxic reactive oxygen species that can selectively damage tumor tissue [20,22]. Porphyrin-based photosensitizers preferentially accumulate in malignant cells, making PDT itself a promising surgical adjunct for treating ovarian micromets [15,23–27]. They have been used in many human clinical trials and naturally accumulate 2–3 fold higher in malignant cells [17,28–30]. Targeting strategies (e.g., EGFR- or folate-directed) have been shown to reduce off-target peritoneal phototoxicity in preclinical models [2,7,15,22,31–36]. Together, these features provide a controlled-release, site-specific drug delivery platform that may improve micromet treatment while minimizing systemic toxicity. Optically triggered combination regimens that integrate imaging and PDT have been positioned as intrinsically theranostic approaches for treatment planning and monitoring [37]. Related imaging-guided phototherapies have leveraged theranostic nanoliposomal photosensitizer formulations and dual-function photosensitizer–fluorophore constructs for fluorescence-guided PDT, supporting the broader “see-and-treat” paradigm for treatment planning and monitoring [38,39].

A primary advantage of this approach is the fluorescence properties of both the PoP carrier and the doxorubicin payload, enabling high-sensitivity, image-guided delivery. PoP fluorescence (~720 nm) can localize nanocarrier accumulation, supporting lesion targeting and treatment planning, while Dox fluorescence (~590 nm) can serve as a readout of drug kinetics and light-triggered release at the target [16,17]. By leveraging a laparoscope equipped with appropriate excitation and emission filtering, these markers can support real-time visualization and quantitative monitoring during minimally invasive CPT.

However, accurately quantifying drug fluorescence concentrations *in vivo* remains challenging due to heterogeneous tissue optical properties and depth-dependent attenuation. Quantitative approaches such as wide-field and laparoscopic/endoscopic spatial frequency domain imaging (SFDI) can map optical properties and support fluorescence correction in scattering tissue [5,40–43]. To bridge the gap between monolayer cell cultures and complex animal models, 3D multicellular spheroid systems provide a critical intermediate platform. Unlike traditional 2D models, spheroids replicate key transport barriers and cellular architecture relevant to peritoneal micrometastases, enabling controlled evaluation of drug distribution, penetration, and fluorescence readouts in a geometry that mimics occult micromet clusters.

In this work, we present a mesoscopic evaluation of light-triggered drug delivery using a custom-designed laparoscopic fluorescence imaging system. This platform was used to monitor uptake and 660-nm-triggered release of long-circulating Dox-PoP (LC-Dox-PoP) liposomes in 3D multicellular spheroid cluster models. By imaging across multiple administered concentrations, we calibrated doxorubicin-associated fluorescence and characterized uptake and release dynamics in spheroids, capturing the transition from the encapsulated state to the therapeutically active diffusive state. Together, these studies establish mesoscopic laparoscopic imaging as a quantitative framework for treatment planning and monitoring of CPT in peritoneal micrometastases. To assess generalizability beyond ovarian cancer, we also evaluate SCC2095sc spheroid clusters, an oral squamous cell carcinoma model relevant to head and neck/oral cancer. Recent head and neck/oral squamous cell carcinoma studies have used 3D spheroid models for quantitative evaluation of image-guided phototherapies, supporting the translational relevance of spheroid platforms for CPT optimization [44–46].

## 2. Results

### 2.1. Fluorescence Calibration in 2D Cell Culture

Fluorescence imaging in 2D culture demonstrated concentration-dependent accumulation of light-activated LC-Dox-PoP within both cell lines. SKOV-3 cells exhibited moderately higher signal intensity than SCC2095sc across matched drug concentrations. As shown in Figure 1e,f, both models exhibited a linear, dose-dependent relationship between administered LC-Dox-PoP concentration and post-activation doxorubicin fluorescence. Signal distribution within wells showed modest heterogeneity (standard deviation < 0.15 across wells), consistent with predominantly intracellular localization rather than uniform surface adsorption.

**Figure 1:**
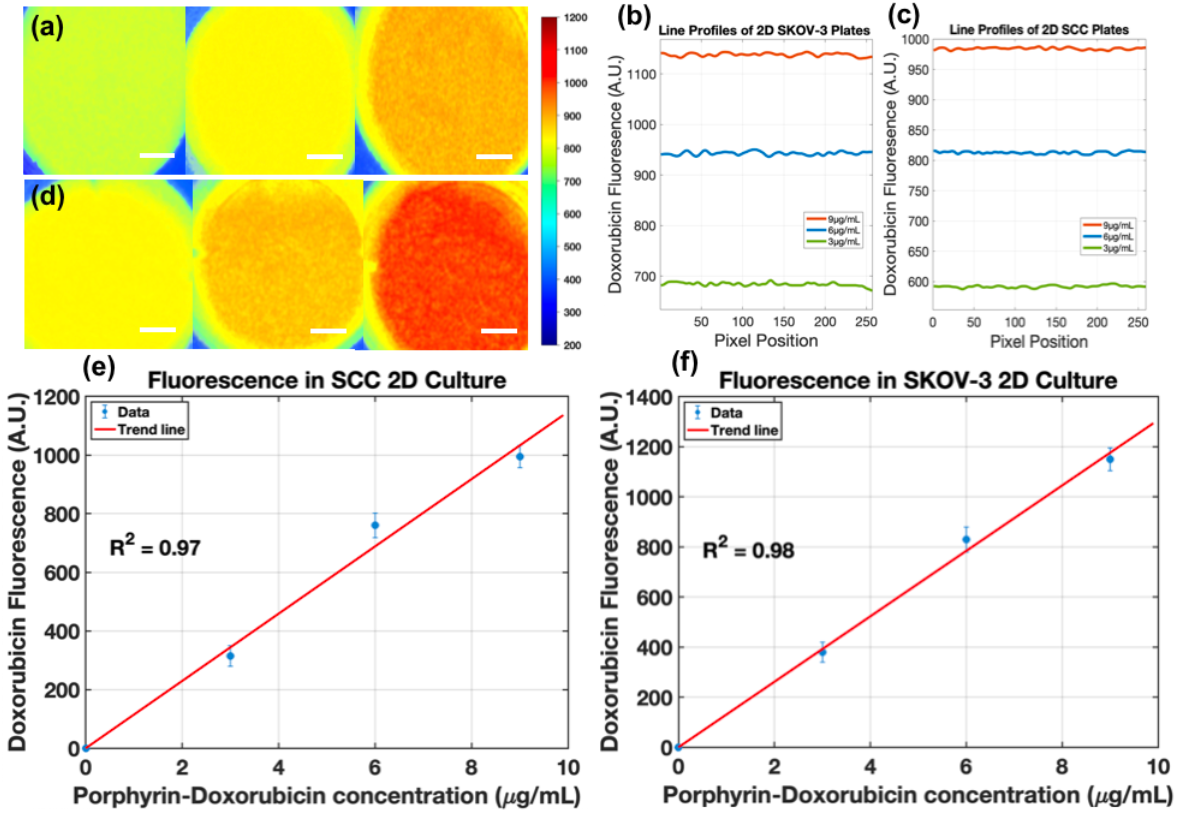
Fluorescent images of SCC2095sc (a) and SKOV-3 (d) cells in 2D culture, showing 3, 6, and 9 µg/mL concentrations of Doxorubicin-Porphyrin liposomes added to the cell media, after 1 min of applied activation light (scale bar = 700µm). (b) Line profile of SCC2095sc and (c) SKOV-3 images obtained at the center of the well plate (e) Linear fit of R^2^ = 0.97 and SKOV-3 fluorescence (f) follows a R^2^ = 0.98 linear trend.

### 2.1. Doxorubicin and Porphyrin Signal in Spheroid Clusters

#### 2.2.1. SCC2095sc Spheroid Clusters

In SCC2095sc spheroid clusters, post-activation doxorubicin fluorescence increased linearly with administered LC-Dox-PoP concentration (1–9 µg/mL), indicating scalable uptake and triggered release without saturation over this range (Figure 2c, R^2^ = 0.98). Representative mesoscopic images (Figure 2a,b) show localized drug-associated fluorescence within spheroid aggregates, with heterogeneous intracluster distribution that is expected in 3D transport-limited geometries. SCC2095sc is a human oral squamous cell carcinoma line, supporting extension of this release-monitoring workflow to head and neck/oral cancer models.

**Figure 2:**
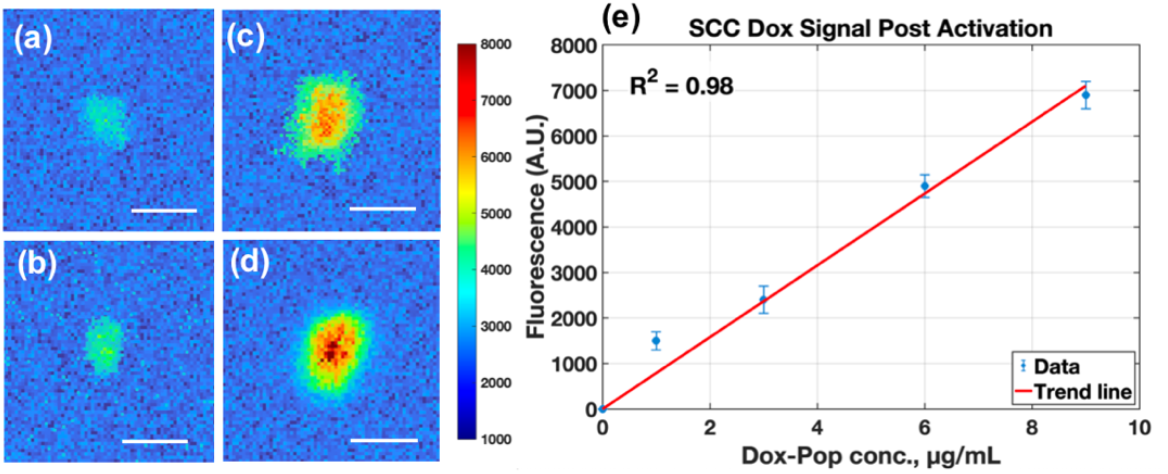
Mesoscopic fluorescent images of SCC cells in 24 well plates, with cultured drug concentration of 1, 3, 6, 9 µg/mL going from left to right. (a) Shows spheroid cluster with full FOV (scale bar = 700µm) and (b) displays digital zoom of fluorescent image of spheroid cluster (scale bar = 300µm). (c) Plots signal increase of spheroid cluster signal within each well with a liner trend (R^2^ = 0.98) with respect to concentration.

Pre-activation porphyrin fluorescence exhibited a linear dose-response relationship, confirming consistent uptake of the liposomal carrier across the tested concentrations (Figure 3e). The porphyrin channel provided higher raw fluorescence counts and clearer localization of spheroid aggregates than the doxorubicin channel under our acquisition settings (compare Figures 2 and 3), consistent with the far-red emission band of PoP (~720 nm) and prior reports of strong porphyrin fluorescence in PoP formulations [10–15,47]. Because the two channels use different excitation/emission bands and acquisition parameters, cross-channel comparisons should be interpreted qualitatively. In this framework, PoP fluorescence primarily supports carrier localization/treatment planning, while doxorubicin fluorescence reports triggered release.

**Figure 3.**
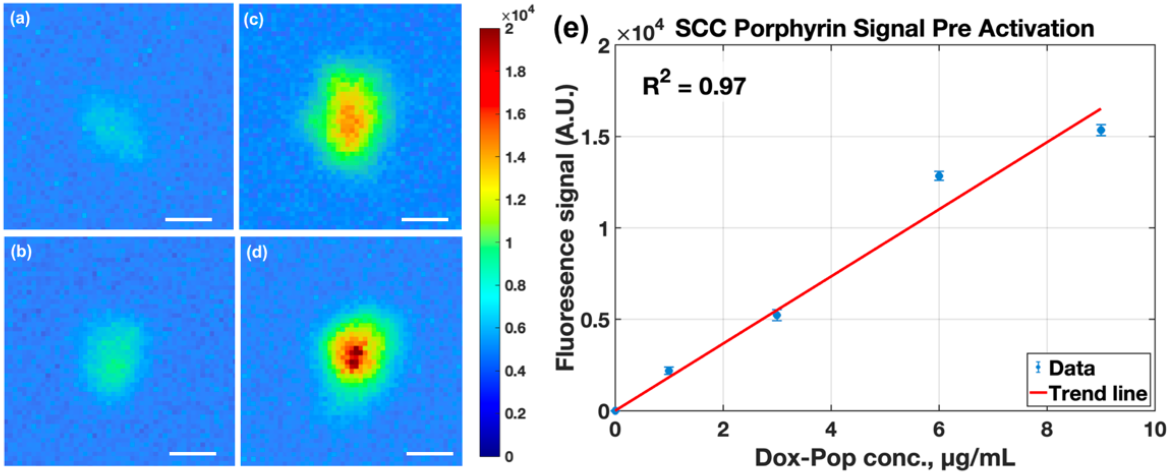
Mesoscopic fluorescence images of pre-activation porphyrin (PoP) signal in SCC2095sc spheroid clusters, with administered LC-Dox-PoP concentrations of 1, 3, 6, and 9 µg/mL in (a-d), displayed as pseudocolor heat maps (common color scale shown) at full field-of-view (scale bar = 700 µm). (e) Mean thresholded porphyrin fluorescence increased linearly with administered concentration (R^2^ = 0.97).

#### 2.2.2. Doxorubicin and Porphyrin Signals in SKOV-3 Spheroid Clusters

In contrast to the compact SCC2095sc spheroids, SKOV-3 clusters exhibited a more dispersed morphology. Despite this structural heterogeneity, post-activation doxorubicin fluorescence increased progressively with administered LC-Dox-PoP concentration (Figure 4b), and digital zoom images highlight non-uniform intracluster drug distribution (Figure 4c). For visualization of small, spatially distributed clusters, the zoomed panels are displayed with locally scaled color maps. Quantitative comparisons were performed on the underlying leakage-corrected fluorescence images using consistent thresholding to preserve linearity across concentrations.

**Figure 4.**
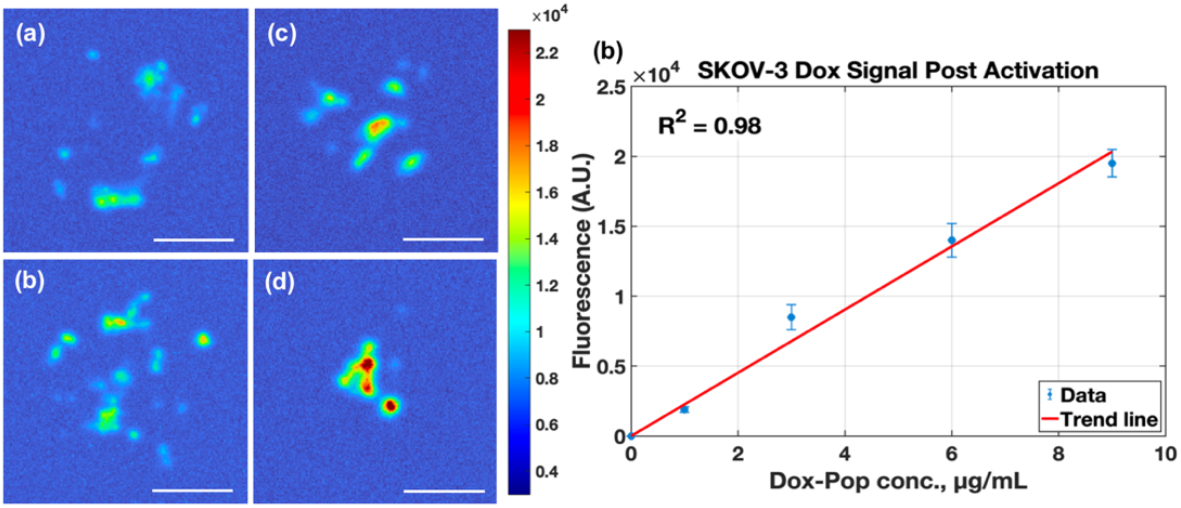
Mesoscopic fluorescence images of post-activation doxorubicin signal in SKOV-3 spheroid clusters after incubation with 1, 3, 6, and 9 µg/mL LC-Dox-PoP (a, left to right). Full field-of-view images are displayed as pseudocolor maps with individual color bars to accommodate the dynamic range across concentrations (scale bar = 700 µm). (b) Mean thresholded doxorubicin fluorescence increased linearly with administered concentration (R^2^ = 0.98). (c) Representative digital zoom images at 3 and 9 µg/mL (scale bar = 300 µm).

In SKOV-3 spheroid clusters, pre-activation porphyrin fluorescence also scaled linearly with administered LC-Dox-PoP concentration (Figure 5e). Under the acquisition settings used here, porphyrin fluorescence exhibited a higher dynamic range than doxorubicin fluorescence (compare Figures 4 and 5), which is consistent with the strong far-red PoP emission and reduced background at longer wavelengths. SKOV-3 clusters also demonstrated higher mean fluorescence than SCC2095sc in both channels, which likely reflects differences in spheroid morphology, uptake, and/or cell density (see Discussion section).

**Figure 5.**
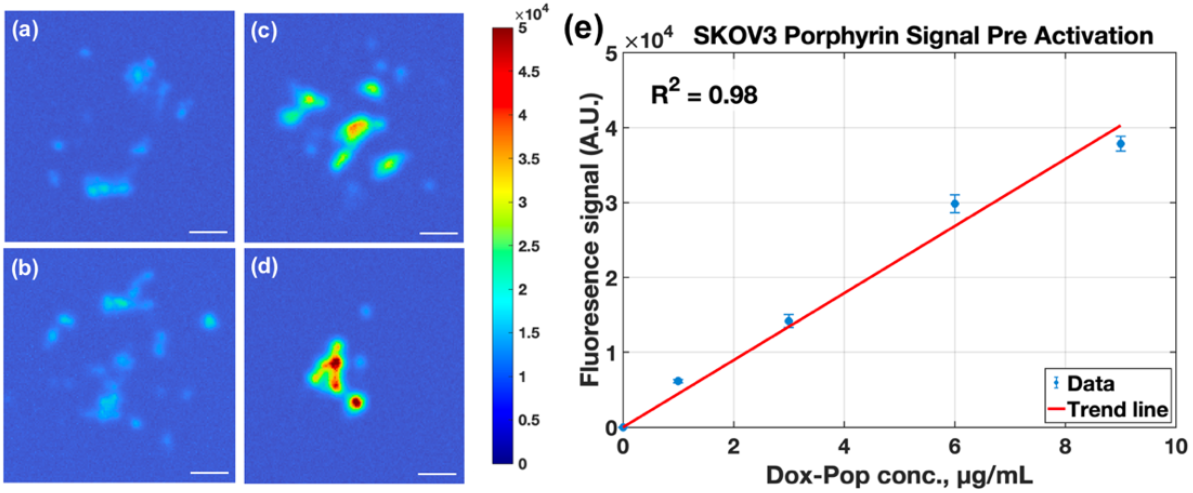
Mesoscopic fluorescence images of pre-activation porphyrin (PoP) signal in SKOV-3 spheroid clusters, with administered LC-Dox-PoP concentrations of 1, 3, 6, and 9 µg/mL in (a-d), displayed as pseudocolor heat maps with a common color scale (scale bar = 700 µm). (e) Mean thresholded porphyrin fluorescence increased linearly with administered concentration (R^2^ = 0.98).

**Figure 6.**
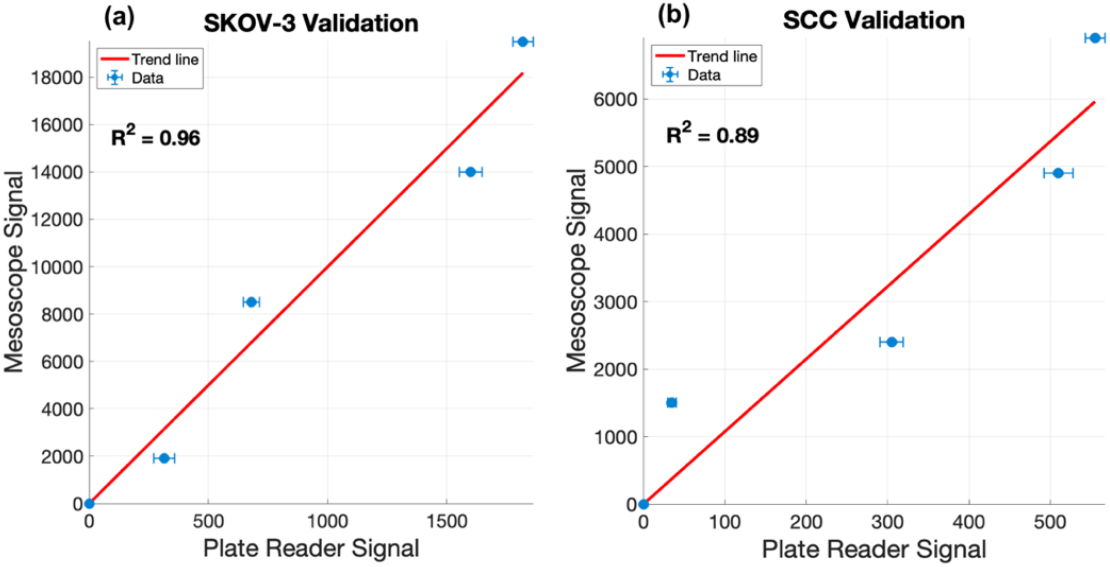
Validation of laparoscopic doxorubicin fluorescence measurements against a standard well-plate reader for SKOV-3 (a) and SCC2095sc (b) spheroid clusters. Linear regression yielded R^2^ = 0.96 (SKOV-3) and R^2^ = 0.89 (SCC2095sc) across the tested LC-Dox-PoP concentrations.

### 2.3. Validation of Fluorescence Signal

To validate the accuracy of the laparoscopic fluorescence workflow, mean doxorubicin fluorescence from each concentration group was compared against standard plate reader measurements. Linear regression demonstrated strong agreement (R^2^ = 0.96 (SKOV-3) and R^2^ = 0.89 (SCC2095sc)) between platforms for both cell models, supporting the use of the laparoscope-derived signal as a quantitative readout of drug-associated fluorescence in the spheroid assay.

### 2.4. Porphyrin Signal in Spheroid Clusters

Across both SKOV-3 and SCC2095sc spheroid clusters, mean porphyrin fluorescence decreased modestly after 660-nm activation light exposure at all tested concentrations (Figure 7), consistent with PoP photobleaching during CPT illumination. This behavior provides complementary feedback such that pre-activation PoP fluorescence can be used to localize nanocarrier accumulation for treatment planning, while a post-illumination decrease in PoP signal can serve as an indicator of delivered light dose/activation, alongside the doxorubicin release readout.

**Figure 7.**
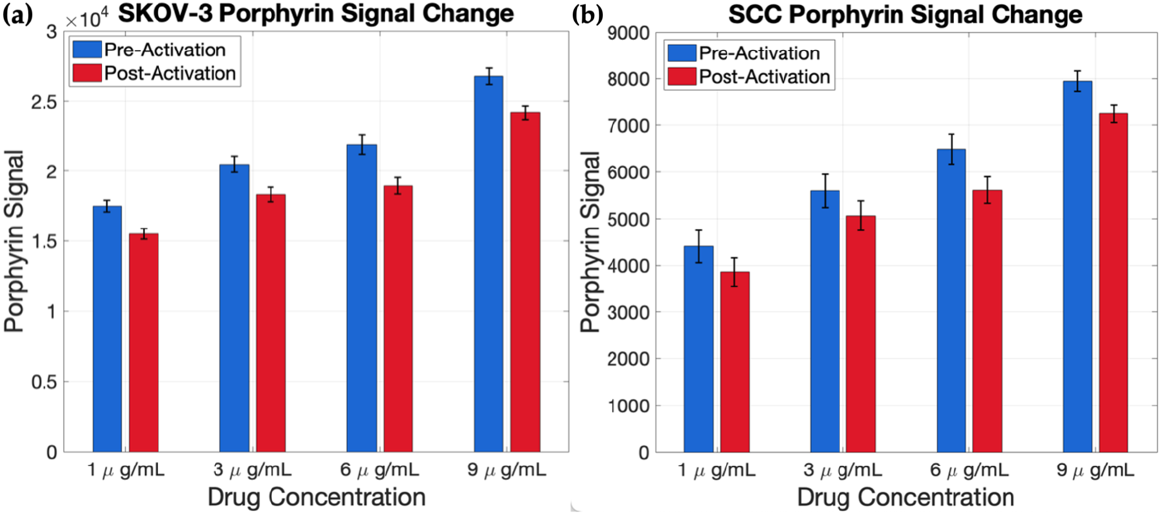
Averaged Porphyrin signal value from SKOV-3 (a) and SCC2095sc (b) spheroids before and after NIR light induced release, showing a slight decrease after light exposure.

Porphyrin signal from the LC-Dox-PoP liposomes was also measured before and after light-triggered release. There was a consistent decrease in porphyrin fluorescence after drug release, suggesting a degree of photobleaching occurring during the light-delivery process. Despite this slight attenuation, the porphyrin signal remained a robust indicator of the initial liposomal carrier distribution throughout the 3D spheroid architecture, with a spatial distribution that closely mirrored the doxorubicin localization within both the SCC and SKOV-3 clusters.

### 2.5. Microscopic Spheroid Clusters

Figure 8 compares spheroid morphology between SKOV-3 and SCC2095sc cell lines 10 days after seeding. SKOV-3 cultures form fewer uniform aggregates with smaller clusters and dispersed cellular material, whereas SCC2095sc forms densely packed, rounded spheroids of varying sizes. These morphological differences likely contribute to the observed differences in fluorescence intensity and spatial distribution between models by altering transport barriers and effective cell density within the imaged clusters.

**Figure 8.**
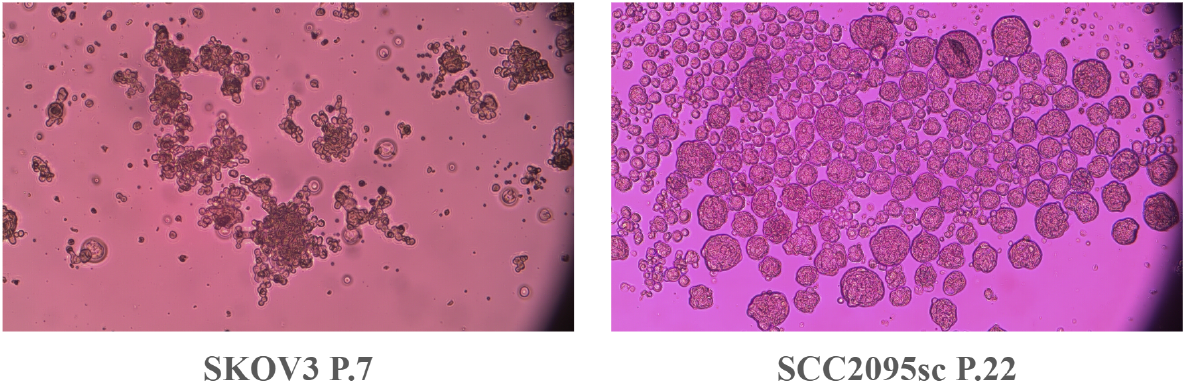
Comparison of spheroid formation in SCC2095sc and SKOV-3 cells 10 days after seeding, demonstrating distinct morphological characteristics. Images were acquired using a 10× objective.

**Figure 9.**
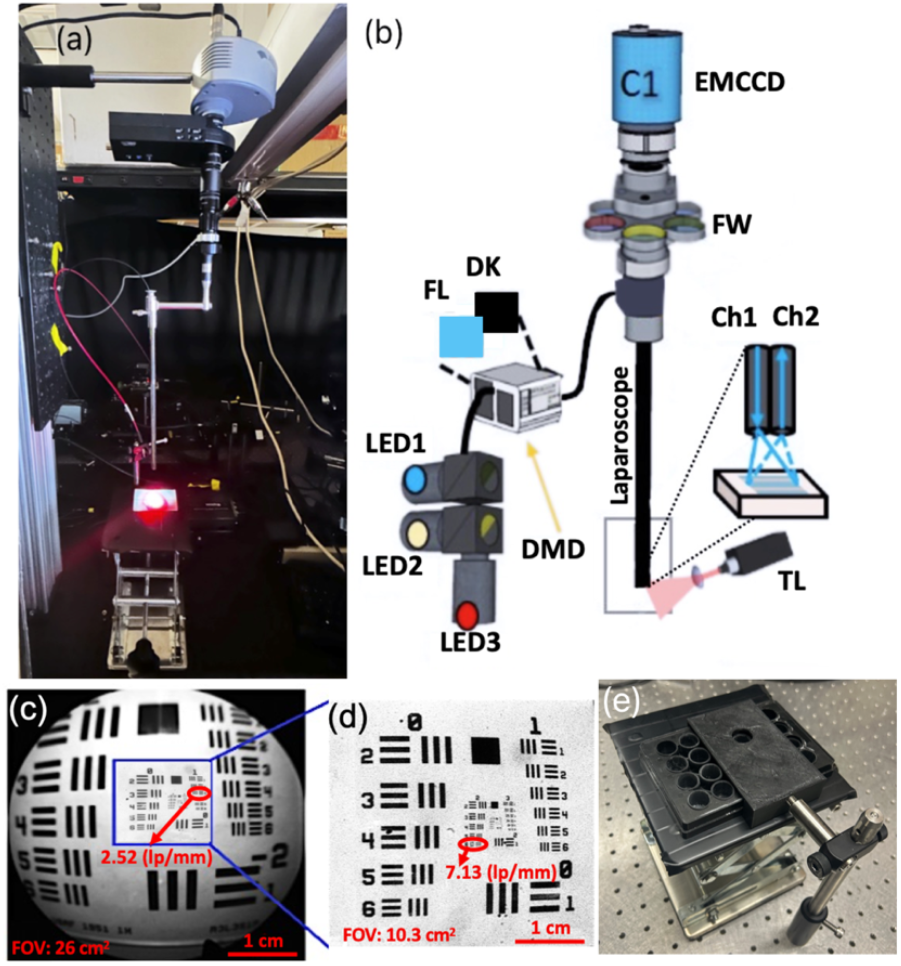
The experimental setup of Doxorubicin release in cell spheroid model, (a) laparoscope SFDI system picture projecting excitation light and the 660 nm treatment light in red, (b) laparoscope SFDI setup design sketch, showing component details (c) Laparoscopic image of fluorescent-positive United States Air Force (USAF) target. (d) Reflectance image of USAF target. Outlined region in (c) provides relative scale of the Fluorescence and Reflectance modalities. (e) Custom well-plate holder with a fixed viewing port and sliding track to ensure consistent spatial alignment of the well plate across sequential imaging sessions.

**Figure 10.**
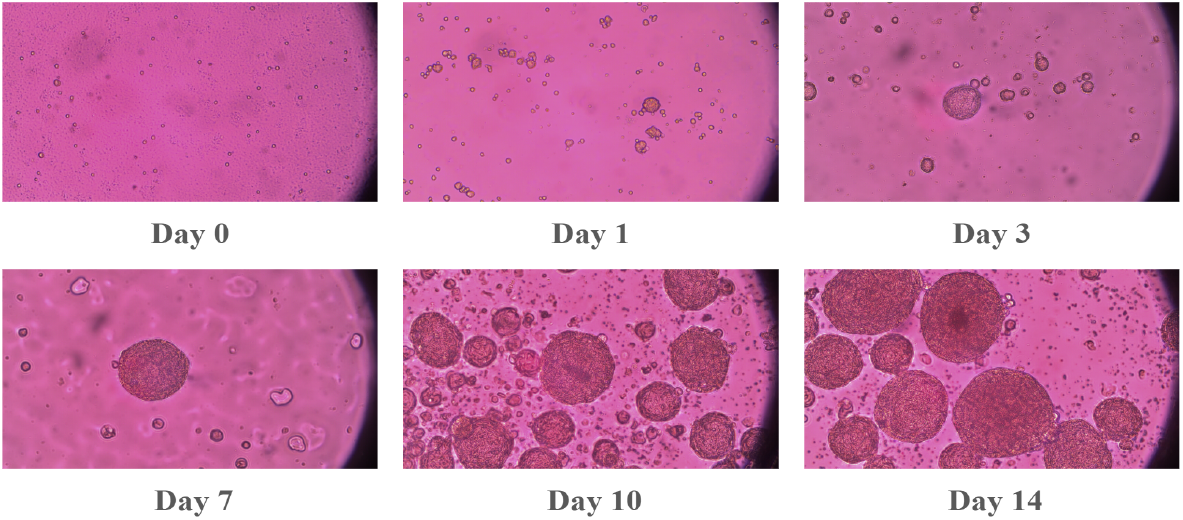
Spheroid formation of SCC2095sc cells. Spheroid formation was monitored at different 8 points (1, 3, 7, 10, and 14 days after seeding the cells). Images were obtained via an inverted microscope using a 20x objective.

**Figure 11.**
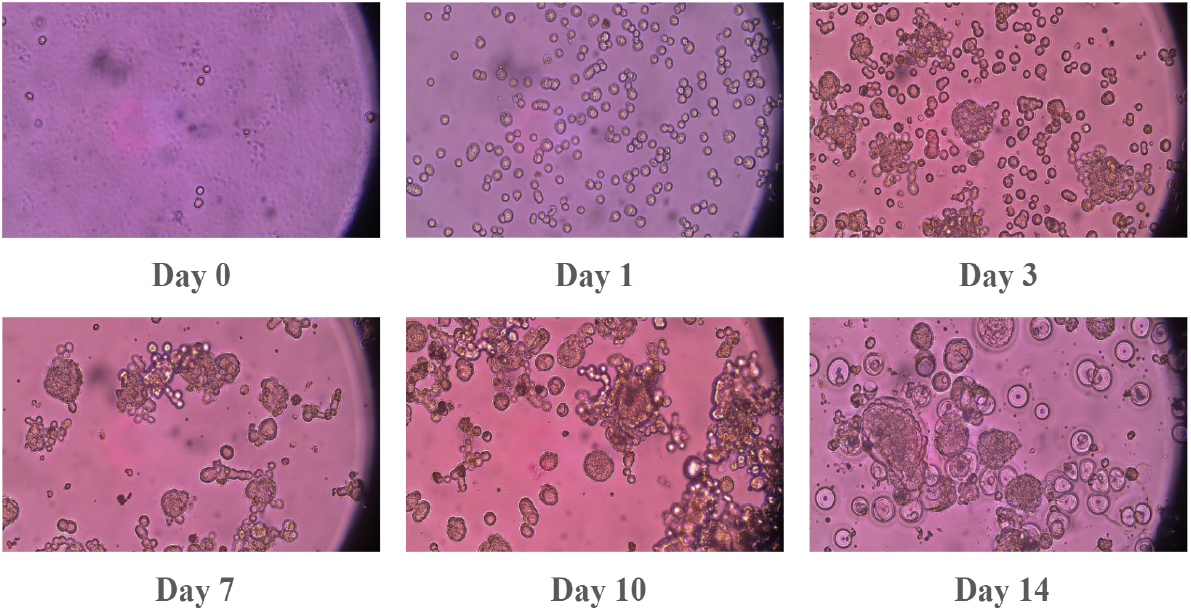
Spheroid formation of SKOV-3 cells. Spheroid formation was monitored at different time points (1, 3, 7, 10, and 14 days after seeding the cells). Images were obtained via an inverted microscope using a 20x objective.

**Figure 12.**
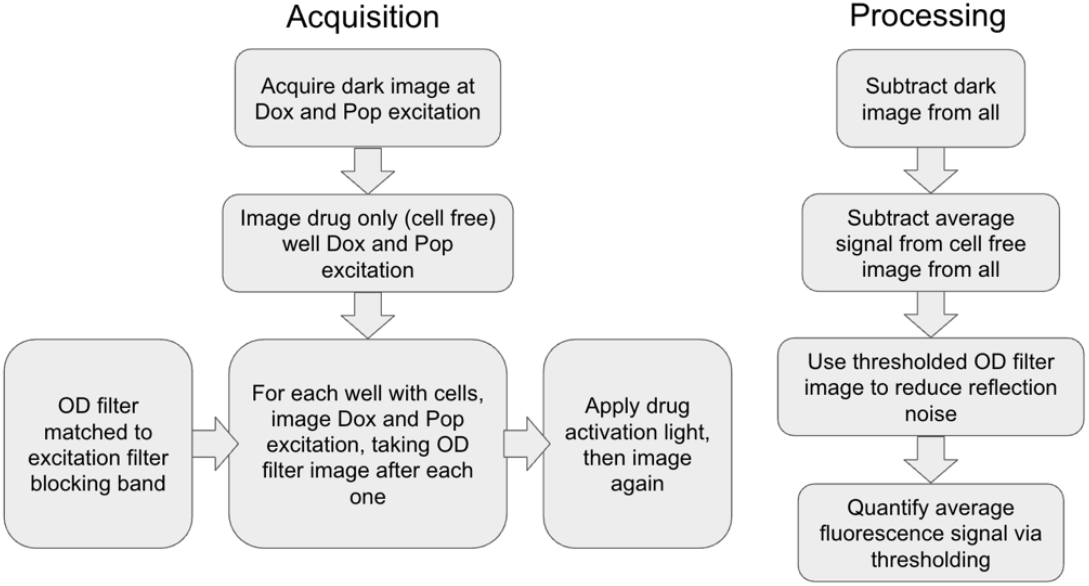
Flowchart of fluorescent imaging and post-processing.

## 3. Discussion

SCC2095sc cells consistently formed dense, uniform, and spherical aggregates characterized by strong cell-cell junctions and a smooth boundary. In contrast, SKOV-3 spheroids exhibited a dispersed morphology with greater variability in size and shape, likely due to reduced expression of adhesion molecules and a more fragmented extracellular matrix. These morphological differences [48] directly influenced the varying fluorescence signals observed between the two cell types. The compact, high-density packing of SCC2095sc spheroids created physical barriers and high interstitial pressure [49] that limited drug diffusion and reduced final signal relative to SKOV-3 spheroids.

Conversely, the porous and irregular architecture of SKOV-3 clusters provided a higher effective surface area and reduced transport resistance, allowing for improved penetration of the liposomes and a higher recorded signal. Furthermore, the similarity in drug uptake profiles between SCC2095sc and SKOV-3 cells in two-dimensional culture suggests that the differences observed in the three-dimensional models are structural rather than purely biochemical.

In this model, the higher fluorescence signal observed in the spheroid center versus the periphery and 2D monolayers is likely an artifact of optical integration. Because widefield mesoscopic imaging projects 3D volumes into 2D images, the signal is integrated along the entire optical pathlength [50–52]. Thus, the thicker center of a spheroid and its stacked cellular layers yield a much higher cumulative intensity than the thinner edges or a single cell layer. Additionally, the central signal concentration likely reflects the 24-hour incubation period, which provides sufficient diffusion time for the necrotic core to act as a metabolic “sink” [53,54]. This represents a key distinction from *in vivo* tumors, where high interstitial pressure, dense matrices and limited blood flow often prevent doxorubicin from reaching the tumor center, highlighting a limitation in how spheroid models replicate complex physiological barriers.

The linear increase in signal as a function of administered LC-Dox-PoP concentration indicates that the uptake and delivery mechanism remains within a linear kinetic range in these assays. Across 1-9 µg/mL range, the system did not reach an apparent saturation point for carrier accumulation and/or drug-associated fluorescence [55], suggesting a predictable and scalable dose-response relationship in three-dimensional tumor models.

The experimental results demonstrated a clear correlation between the fluorescence signal and the concentration of the delivered therapeutic. In both the 2D monolayer and the 3D spheroid models [56], a proportional relationship was observed in the signals recorded from the samples. As the concentration of the drug increased, the fluorescence intensity scaled accordingly. While these initial findings suggest that the fluorescence signal may serve as a proxy for drug concentration and tumor volume, further studies involving a broader range of concentrations are required to establish a robust linear relationship and a definitive calibration curve [22]. The use of light-triggered Doxorubicin in conjunction with a filtered laparoscopic system provides a robust framework for fluorescence-guided surgery. Additionally, the spheroid model here provides a more robust estimate of in vivo drug kinetics due to the more realistic properties of the tumor microenvironment [28–30]. By successfully isolating the signals of Dox in light triggered liposomes and subtracting background artifacts via ND filtering, we have shown that it is possible to visualize simulated micromet cell clusters in an in vitro model.

Importantly, the strong linearity of the fluorescence dose–response across both 2D and 3D models (R^2^ ≥ 0.97) and the cross-platform agreement with plate reader measurements (R^2^ = 0.89–0.96) indicate that the laparoscopic workflow can provide a quantitative readout of nanocarrier delivery and light-triggered release. In the context of chemophototherapy, such fluorescence-based feedback could enable lesion-specific treatment planning (carrier localization via PoP) and real-time monitoring of on-target drug activation (Dox release), supporting optimization of light dosimetry while reducing off-target exposure. Moreover, the higher signal levels observed in the porphyrin channel relative to doxorubicin under our imaging conditions suggest that PoP fluorescence may be more robust for intraoperative localization and treatment planning, whereas doxorubicin fluorescence remains valuable as a release-specific reporter but may be more susceptible to attenuation and background at shorter wavelengths. This concept is consistent with prior demonstrations of imaging-guided PDT using porphyrin-based nanoliposomal platforms, where fluorescence readouts were used to localize therapeutic accumulation and guide light delivery[38]. Moreover, time-gated fluorescence tomography has been used to image PDT photosensitizer distributions in vivo, providing a route toward depth-sensitive intensity/lifetime quantification in future extensions of this workflow [57].

Beyond peritoneal disease, the inclusion of an oral squamous cell carcinoma model (SCC2095sc) highlights the potential applicability of this quantitative fluorescence workflow to head and neck/oral cancer. In head and neck surgery, local recurrence is often driven by microscopic residual disease; a locally activated, fluorescence-monitored chemophototherapy strategy could complement postoperative adjuvant PDT or chemotherapy by confining activation to the surgical bed while limiting off-target exposure. Future studies will translate these in vitro findings to orthotopic head and neck models and evaluate treatment-response monitoring under clinically relevant illumination geometries. Consistent with this rationale, Mallidi and colleagues have evaluated intraoral PDT light-delivery systems for oral lesions and have reported head and neck/oral cancer 3D spheroid models for quantitative evaluation of photoactive agents and image-guided phototherapies [44–46,58]. In addition, studies on HPPH-based porphyrin analogs have shown that peripheral substitutions (e.g., charge/lipophilicity) can substantially influence cellular uptake/retention and PDT efficacy, motivating quantitative fluorescence readouts for selecting and optimizing photoactive agents for specific tumor contexts [59]. Additionally, photoactivated HPPH-liposomes have demonstrated tumor-selective uptake and phototherapy-induced tumor control in chemoradioresistant head and neck cancer patient-derived xenograft models, supporting translation of porphyrin-liposomal photoactivated delivery strategies [60]. Clinical studies of HPPH-mediated photodynamic therapy in oral cavity/head and neck cancer further support the feasibility of porphyrin-based, locally activated adjuvant approaches [61–71].

## 4. Materials and Methods

### 4.1. Laparoscopic SFDI System

The laparoscopic SFDI system was constructed and validated earlier [72] as shown in Figure 7. Figure 7a shows the sample imaging setup for release experiment, and figure 7b details the projection with digital micromirror device (DMD) and imaging components within the laparoscopic system.

In our laparoscopic fluorescence imaging system, illumination was provided by three high-power LEDs at 490 nm, 590 nm, and 656 nm, which were coupled to the DMD via a liquid light guide (Mightex, Ontario, Canada). Both the LEDs and DMD were remotely controlled using MATLAB. Excitation light was projected through a 2.4 mm imaging fiber containing 13,000 elements (Asahi Kasei, Tokyo, Japan), which was coupled to a fixed laparoscope and delivered light to the distal end facing the tissue. A custom objective lens at the fiber tip collimated the output and provided uniform illumination across the tissue surface. The excitation signal were collected through the laparoscope optics, including the laparoscope (8912.43, R. Wolf, Vernon Hills, IL, USA), a zoom coupler (Accu-Beam, TTI Medical, CA, USA), two 25 mm achromatic lenses, a filter wheel, and an aperture (ThorLabs, NJ, USA), before being detected by an EMCCD camera (1004 × 1002 pixels, Luca, Andor, Belfast, Ireland). The camera was focused to capture signal over a 3.2 × 3.2 cm field of view (FOV). At a 5.1 × 5.1 cm FOV, the system could resolve group 1, element 3 of the target, corresponding to a line width of 198.43 µm, which is equivalent to 2.52 line pairs/mm as shown in Figure 7c. Figure 7d demonstrates a higher magnification (with a smaller FOV of 3.2 cm × 3.2 cm), at which the endoscopic system can resolve group 2, element 6 of the target, corresponding to line width 70.15 µm, which is equivalent to 7.13 line pairs/mm.

A 490 nm LED with a 490 ± 10 nm bandpass filter was used to excite Dox fluorescence, and the 656 nm LED was used to excite Porphyrin fluorescence. To isolate the fluorescent signal of doxorubicin, a 530 nm long-pass filter and a 593 ± 40 nm fluorescent filter were used. Neutral density (ND) filters, matched to the transmission of the fluorescent filters, are used to obtain reflectance data and remove specular reflections and signal leakages, as described further in 4.4. To capture fluorescent signal from the porphyrin loaded liposomes, a 750 ± 50nm fluorescent filter was used with the 656nm excitation light. A custom 3D-printed black structure (Figure 7e) serves as a physical stop for multi-well plates. This ensures precise alignment and a constant imaged region across successive captures and filter changes while mitigating spectral reflections.

### 4.2 Preparation of long-circulating Dox in PoP Liposomes

The details of our PoP Dox liposome formulation have been described in [1,4,5]. PoP-liposomes incorporated PoP photosensitizer in a recent study for the development of improved peptide-based cancer vaccines, underscoring the versatility of the drug delivery platform [13]. Briefly, PoP-liposomes were synthesized from pyro-lipid through esterification of pyro with lyso-C16-PC, using 1-Ethyl-3-(3-dimethylaminopropyl) carbodiimide (EDC) and 4-dimethylaminopyridine (DMAP) in chloroform. The liposomes were formed by dispersing Porphyrin-lipid, PEGylated-lipid, cholesterol, and distearoylphosphatidylcholine in chloroform, followed by solvent evaporation. A 20 mg/mL lipid solution was extruded through a high-pressure lipid extruder with a 250 mM ammonium sulfate solution using polycarbonate membranes of 0.2, 0.1, and 0.08 µm pore size, sequentially stacked and passed through the extruder 10 times. Free ammonium sulfate was removed by overnight dialysis in a 10% sucrose solution with 10 mM HEPES at pH 7. Dox was loaded by incubating the liposomes at 60°C for 1 hour, achieving a loading efficacy of over 90% as confirmed by G-75 column tests. The self-assembly status and elution position of PoP-liposomes were tracked using 420 nm excitation and 670 nm emission, while Dox was detected using 480 nm excitation and 590 nm emission in a fluorescence plate reader (TECAN Safire).

### 4.3 Spheroid Model

Human epithelial ovarian adenocarcinoma cell lines SKOV-3 were purchased from the American Type Culture Collection (ATCC, HTB-77, Manassas, VA, USA) and were maintained in McCoy’s 5A medium supplemented with 10% Fetal Bovine Serum (FBS) and 1% antibiotic-antimycotic. Human oral squamous cell carcinoma (OSCC) SCC2095sc (gift from Professor S. Mallery, Ohio State University) were cultured in Advanced DMEM-Serum Reduced Medium supplemented with 1% L-Glutamine, 5% heat-inactivated FBS, and 1% antibiotic-antimycotic. Media and supplements were purchased from Gibco (Grand Island, NY, USA). Cells were cultured at 37°C with 5% CO^2^ in a humidified incubator. Upon reaching 80–90% confluence, cells were harvested using 0.25% trypsin-EDTA and passaged at a 1:3 ratio. To facilitate three-dimensional (3D) spheroid formation, 12-well plates were coated with Poly (2-hydroxyethyl methacrylate) (Poly-HEMA; Sigma-Aldrich) to create non-adherent surfaces. A 1.2% (w/v) Poly-HEMA working solution in 95% ethanol was applied to each well (240 µL/well) and allowed to dry for 24–48 hours at room temperature. Prior to seeding, plates were sterilized via UV exposure for 30 minutes and rinsed twice with PBS and once with Hank’s balanced salt solution (HBSS). Spheroids were generated by seeding 10,000 viable cells per well into the Poly-HEMA-coated plates. The final volume in each well was adjusted to 1 mL with complete growth media. Spheroid morphology, growth, and integrity were monitored longitudinally (from Day 0 to Day 15) using an inverted microscope. Qualitative assessments focused on spheroid shape, compactness, and the presence of cellular debris.

Chemophototherapy was performed using a porphyrin-phospholipid liposomal formulation of doxorubicin (LC-Dox-PoP) described in 4.4. Experiments were initiated once spheroids reached maturity, typically 14–16 days post-seeding for SKOV-3 and 10– 12 days for SCC2095sc. Spheroids were treated with LC-Dox-PoP at final concentrations of 1, 3, 6, and 9 µg/mL. Control groups received complete growth medium without the drug and “blank” well plates with drug concentrations but no cell presence were used for fluorescence signal calculation (see 4.4). To ensure uniform drug distribution, a designated volume of medium was replaced or supplemented with the LC-Dox-PoP working solution to reach the target concentrations. Following drug administration, plates were protected from light and incubated for 24 hours. Fluorescence imaging was performed prior to and immediately after light triggered release of LC-Dox-PoP liposomes via exposure to a 650 nm laser of 350 *mW/cm*^2^ fluence for 1 minute per well. After mesoscopic imaging, well plates were rinsed three times with PBS and filled with phenol-red free media prior to well plate imaging at Ex = 480 nm, Em = 590 nm (TECAN Infinite M Plex).

### 4.4 Fluorescent Imaging and Image Processing

Prior to imaging the spheroid clusters, calibration images of dark signal, drug filled wells without cells and water filled wells are taken at all filters to account for sensor noise, extracellular drug fluorescence and leakages respectively. For each experimental well, the doxorubicin and porphyrin signals were captured by illuminating the samples at their respective excitation peaks using the appropriate fluorescent and ND filters. Following this initial characterization, activation light was applied to trigger drug release, after which the fluorescent signals were re-imaged to quantify the change in localized drug concentration.

To correct for excitation light leakage into the fluorescence detection channel, an excitation-only reference image was acquired immediately prior to each fluorescence image under identical excitation conditions using a neutral-density (ND) filter with the same optical density (OD) value as the respective porphyrin and doxorubicin fluorescent filters. After dark subtraction, the ND image is assumed to be a linearly scaled version of leakages in the image taken with the fluorescent filters with the same OD blocking filters. To derive this scaling factor, calibration images without fluorophores (water filled wells of identical volume to cell wells) were taken prior to fluorescence imaging at all filter configurations, where the true fluorescence signal is 0. We then perform an Ordinary Least Squares regression over these images to determine our scaling factor α:

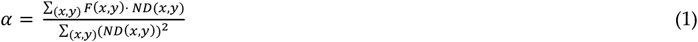

Where (x,y) is the set of pixel coordinates for the ND and fluorescent (F) images. To prevent overcorrection, a spatially weighted mask (*M*) is derived from the normalized signal from the ND filter. Then, the weighted and scaled leakage estimate is subtracted from the processed fluorescence image:

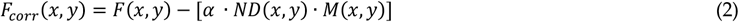

This method corrects for leakage at the cell level, which would be standard across the well plate, and leakage due to reflections caused by the plastic well plate material that could confound signal. This approach preserves linearity, accounts for spatially localized leakage, and minimizes subtraction of true fluorescence signal.

Final quantification of the drug signal was achieved through segmentation, defined by the selection of pixels with intensity values exceeding a threshold that corresponded to the background fluorescence of the culture media. This segmentation process ensured that the average signal intensity reflected only the drug accumulated within the cellular structures, providing a representation of the uptake and release kinetics within the spheroid clusters specifically.

## 5. Conclusions

This study demonstrates a filtered laparoscopic fluorescence imaging workflow for quantitative monitoring of light-triggered LC-Dox-PoP delivery in 2D cultures and 3D spheroid clusters. Dox fluorescence scaled linearly with administered concentration (R^2^ = 0.97-0.98 in 2D and R^2^ = 0.98 in spheroid clusters) and agreed with standard plate reader measurements (R^2^ = 0.89-0.96). Complementary PoP fluorescence supported localization of nanocarrier distribution and remained detectable after activation despite modest photobleaching. Together, these results support dual-fluorescence PoP liposomes as a theranostic platform and establish quantitative imaging readouts that could guide lesion-specific light dosimetry and monitor treatment response during intraoperative chemophototherapy for peritoneal micrometastases. Future work will evaluate in vivo performance and incorporate optical-property correction strategies to improve quantitative robustness in heterogeneous tissue. Demonstrating performance in both SKOV-3 ovarian and SCC2095sc oral squamous cell carcinoma spheroids supports broader translation of fluorescence-guided chemophototherapy monitoring to additional surgical adjuvant settings, including head and neck/oral cancer [60,65,70].

## Acknowledgments

The authors acknowledge the funding support from NIH/NCI R01 (5R01CA243164-06). We also thank Professor Susan R. Mallery (The Ohio State University) for providing SCC20905sc cells.

## Author Contributions

U.S. conceived and designed the experiments; R.A. designed the device, E.K, SAS, B.A. performed the experiments; E.K. analyzed the data; R.A., U.S., E.K., SAS wrote the paper.

## Institutional Review Board Statement

“Not applicable.”

## Informed Consent Statement

“Not applicable.”

## Data Availability Statement

The data that support the findings of this study are available from the corresponding author upon reasonable request.

